# Deep learning reveals evolutionary conservation and divergence of sequence properties underlying gene regulatory enhancers across mammals

**DOI:** 10.1101/110676

**Authors:** Ling Chen, Alexandra E. Fish, John A. Capra

## Abstract

In mammals, genomic regions with enhancer activity turnover rapidly; in contrast, gene expression patterns and transcription factor binding preferences are largely conserved. Based on this conservation, we hypothesized that enhancers active in different mammals would exhibit conserved sequence patterns in spite of their different genomic locations. We tested this hypothesis by quantifying the conservation of sequence patterns underlying histone-mark defined enhancers across six diverse mammals in two machine learning frameworks. We first trained support vector machine (SVM) classifiers based on the frequency spectrum of short DNA sequence patterns. These classifiers accurately identified many adult liver, developing limb, and developing brain enhancers in each species. Then, we applied these classifiers across species and found that classifiers trained in one species and tested in another performed nearly as well as classifiers trained and tested on the same species. This indicates that the short sequence patterns predictive of enhancers are largely conserved. We also observed similar cross-species conservation when applying the models to human and mouse enhancers validated in transgenic assays. The sequence patterns most predictive of enhancers in each species matched the binding motifs for a common set of TFs enriched for expression in relevant tissues, supporting the biological relevance of the learned features. To test the conservation of more complex sequences patterns, we trained convolutional neural networks (CNNs) on enhancer sequences in each species. The CNNs demonstrated better performance overall, but worse cross-species generalization than SVMs, suggesting the importance of combinatorial interactions between motifs, but less conservation of these more complex sequence patterns. Thus, despite the rapid change of active enhancer locations between mammals, cross-species enhancer prediction is often possible. Furthermore, short sequence patterns encoding enhancer activity have been maintained across more than 180 million years of mammalian evolution, with evolutionary change in more complex sequence patterns.

**Author summary:** Alterations in gene expression levels are a driving force of both speciation and complex disease; therefore, it is of great importance to understand the mechanisms underlying the evolution and function gene regulatory DNA sequences. Recent studies have revealed that while gene expression patterns and transcription factor binding preferences are broadly conserved across diverse animals, there is extensive turnover in distal gene regulatory regions, called enhancers, between closely related species. We investigate this seeming incongruence by analyzing genome-wide enhancer datasets from six diverse mammalian species. We trained two machine-learning classifiers—a *k*-mer spectrum support vector machine (SVM) and convolutional neural network (CNN)—to distinguish enhancers from the genomic background. The *k*-mer spectrum SVM models the occurrences of short sequence patterns while the CNN models both the short sequences patterns and their combinatorial patterns. Both the SVM and CNN enhancer prediction models trained in one species are able to predict enhancers in the same cellular context in other species. However, CNNs performed better at predicting enhancers in each species, but they generalize less well across species than the SVMs. This argues that the short sequence properties encoding regulatory activity are remarkably conserved across more than 180 million years of mammalian evolution with more evolutionary turnover in the more complex combinations of the conserved short sequence motifs.

## Introduction

Enhancers are genomic regions distal to promoters that bind transcription factors (TFs) to regulate the dynamic spatiotemporal patterns of gene expression required for proper differentiation and development of multi-cellular organisms [1,2]. It is critical to understand the mechanisms underlying enhancer evolution and function, as alterations in their activity influence both speciation and disease [3–5]. Recent genome-wide profiling of TF occupancy and histone modifications associated with enhancer activity revealed that the regulatory landscape changes dramatically between species—both enhancer activity and TF occupancy at orthologous regions distal to promoters are extremely variable across closely related mammals [6–12]. However, the gene regulatory circuits [13] and expression of orthologous genes in similar tissues are largely conserved across mammals [14–16]. Much of the gene regulatory machinery is also conserved; TFs and the short DNA motifs they bind are highly similar between human, mouse, and fly [17–20]. In short, there is considerable change in the enhancer activity of orthologous regions across mammals, despite the relative conservation of gene expression and TF binding preferences.

The rapid turnover in enhancer activity between orthologous regions in different species has largely been attributed to differences in the DNA sequences of the elements involved, rather than differences in the broader nuclear context [21–25]. Genome-wide profiles of TF binding have shown that 60–85% of binding differences in human, mouse, and dog for the TFs CEBPα and HNF4α can be explained by genetic variation that disrupts their binding motifs [23]. Genetic differences are also often responsible for differential enhancer activity between more closely related species; for example, variation in TF motifs at orthologous enhancers was predictive of activity differences between human and chimp neural crest enhancers [25]. This suggests that, while there is turnover at orthologous sequences, sequence properties predictive of enhancer activity may still be conserved.

Until recently, investigation of the conservation of enhancer sequence properties across mammalian evolution has been hampered by a lack of known enhancers across diverse species within the same cellular context. The canonical definition of enhancer activity is the ability to drive expression in transgenic reporter assays [1,26], which cannot currently be scaled to assess regulatory potential genomewide. However, high-throughput assays such as ChIP-seq can assess histone modifications associated with enhancer activity [27,28] to identify putative enhancers genome-wide in many tissues and species [12,29]. Using known enhancers, machine learning approaches have learned their sequence properties and successfully distinguished enhancers active in specific cellular contexts from both the genomic background and enhancers active in other tissues [30–39]. Moreover, some of these studies suggested the potential for cross-species enhancer prediction. For instance, the similarity of co-occurrence of sequence patterns can be used to identify orthologous enhancers in distantly related Drosophila species [40]. Furthermore, annotated cis-regulatory modules (CRMs) in Drosophila can predict CRMs in highly diverged insect species based on binding site composition similarity [41]. However, TF binding sites appear to evolve and turnover much more rapidly between closely related mammals than Drosophila species [10,42]. In mammals, a machine learning model trained with mouse enhancers accurately predict orthologous regions of the human genome [31]; however, the rapid turnover of enhancer activity between human and mouse suggests that the majority of these orthologous regions are not human enhancers [12]. These previous studies suggest the potential for evolutionary conservation of sequence properties of mammalian enhancers, but comprehensive genome-wide quantification of the degree and dynamics of this conservation is needed.

In this study, we investigate the degree of regulatory sequence property conservation by applying machine learning classifiers to genome-wide enhancer datasets across diverse mammals. We first confirm that Support Vector Machine (SVM) classifiers trained using short DNA sequence patterns can accurately identify many enhancers genome-wide in the adult liver [12], developing limb and developing brain [29]. Then, by using classifiers trained in one species to predict enhancers in the others, we demonstrate that enhancer sequence properties are conserved across species, even though the enhancer activity of specific loci is not. We establish the robustness of this conservation to different enhancer identification techniques by showing that classifiers trained using high-confidence human and mouse enhancer sequences validated in transgenic assays also generalize across species, and are similar to classifiers trained on histone-modification-defined enhancers. Furthermore, the short DNA patterns most predictive of enhancer activity in each species matched a common set of binding motifs for TFs enriched for expression in relevant tissues. This suggests the patterns learned by classifiers capture biologically relevant sequences that influence TF binding. In addition to SVM classifiers, we also trained convolutional neural networks (CNNs) on enhancers in each species. The multilayer structures of CNNs can learn predictive short DNA motifs as well as combinations of motifs at different levels of complexity, and therefore are promising for modeling the complex interactions between TFs [36–39,43–45]. The CNNs predicted enhancers with higher accuracy than *k*-mer SVM models, but the CNNs generalized less well across species, suggesting less conservation of more complex sequence patterns. Together, our results argue that, though there is rapid change of active gene regulatory sequences between mammalian species, the short sequence patterns of the enhancer regions encoding regulatory activity have been conserved over 180 million years of mammalian evolution. Furthermore, the combinatorial rules combining these short sequence patterns may be more divergent between species. Our findings also suggest avenues for improved enhancer identification within and between species and establish a framework for future exploration of the conservation and divergence of regulatory sequence properties between species.

## Results

### Enhancers can be predicted from short DNA sequence patterns in mammals

Genome-wide enhancer activity across many mammalian species was recently assayed by profiling enhancer-associated histone modifications in the adult liver [12], developing limb [8] and developing brain [46]. Certain chemical modifications to histones, such as acetylation of lysine 27 of histone H3 (H3K27ac) and lack of trimethylation of lysine 4 of H3 (H3K4me3), are significantly associated with active enhancers. Determining the genomic locations of these modifications via ChIP-seq provides a genome-wide proxy for the active enhancer landscape [27,28]. For brevity, we refer to genomic regions with enhancer-associated histone modification combinations identified in these previous studies as “enhancers.”

For each species and tissue, we evaluated how well short DNA sequence patterns identified enhancers. We quantified DNA sequence patterns present in each genomic region by computing its *k*-mer spectrum—the observed frequencies of all possible nucleotide substrings of length *k*. We then trained SVM classifiers on the *k*-mer spectra to distinguish enhancers from random genomic regions matched to the enhancers on various attributes, such as length, GC-content, and repeat-content, as appropriate. We trained and evaluated our classifiers on both unbalanced and balanced positive and negative sets (see CNN results and Methods). The unbalanced set contains ten times as many negative non-enhancer regions as enhancers to reflect the fact that most of the genome does not have enhancer activity; in this test, we trained with different misclassification costs for positives and negatives (Methods). We report the unbalanced results in this section and the balanced results in the comparisons with CNN models below. We used ten-fold cross validation to evaluate classifier performance. We quantified performance by computing the average area under receiver operating characteristic (auROC) and precision-recall (auPR) curves over the ten cross-validation runs (Figure 1; Methods).

**Fig 1.**
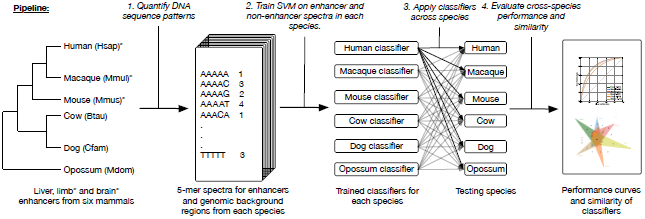
Overview of the framework for evaluating DNA patterns predictive of enhancer activity across diverse mammals. Starting with liver, limb and brain enhancers and genomic background regions from six mammals, the first step of the pipeline quantified each of these genomic regions by their 5-mer spectrum—the frequency of occurrence of all possible length five DNA sequence patterns. Using the spectra as features, we trained a spectrum kernel support vector machine (SVM) to distinguish enhancers from non-enhancers in each species and evaluated their performance with ten-fold cross validation. Then, we applied classifiers trained on one species to predict enhancer activity in all other species. Finally, we evaluated the performance of cross-species prediction compared to within species prediction and compared the most predictive features in classifiers from different species. Limb and brain enhancer data were only available for human, macaque, and mouse.

We first evaluated the ability of classifiers trained on 5-mer spectra to identify liver enhancers in six representative mammals: human, macaque, mouse, cow, dog and opossum. These species were selected as representatives, since they each come from a different clade and have high-quality genome builds. As expected from previous work [31,32,47], all classifiers could distinguish active liver enhancers from length-matched background regions; auROCs ranged from 0.78 in dog to 0.84 in mouse (Figure 2a, PR curves in Figure S1a). Next, we trained 5-mer spectrum SVM classifiers to predict enhancers active in limb and brain for human, macaque, and mouse. Again, classifiers accurately distinguished enhancers from the background with even stronger performance than the liver classifiers. The limb classifiers achieved auROCs of ∽0.89 in each species (Figure 2b; PR curves in Figure S1b), and the brain classifiers had auROCs from 0.90–0.93 (Figure 2c; PR curves in Figure S1c).

**Fig 2.**
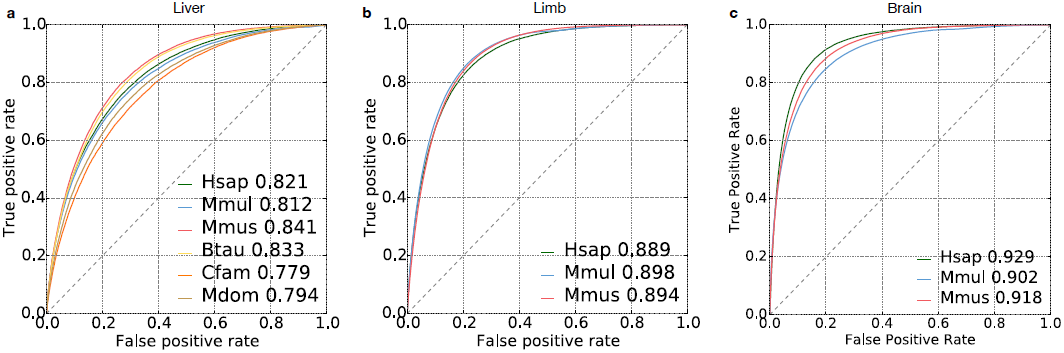
Performance of DNA sequence-based enhancer identification in diverse mammals. (a) ROC curves for classification of liver enhancers vs. the genomic background in six diverse mammals: human (Hsap), macaque (Mmul), mouse (Mmus), cow (Btau), dog (Cfam), and opossum (Mdom). (b) ROC curves for classification of developing limb enhancers in human, macaque, and mouse. (c) ROC curves for classification of developing brain enhancers in human, macaque, and mouse. Area under the curve (AUC) values are given after the species name. Tenfold cross validation was used to generate all ROC and PR curves (Figure S1a,b, c).

The choice of *k* did not significantly influence performance; the auROCs for human liver classifiers are 0.81, 0.82, 0.82, 0.82, respectively across a *k* of 4, 5, 6, and 7. We also explored the application of classifiers based on more flexible *k*-mer features, i.e., the gappy and mismatch *k*-mer kernels [48], but they did not significantly improve performance (Figure S2; Methods). These results illustrate that SVMs trained only on DNA sequence patterns can distinguish many enhancers from background sequences across a variety of mammals for three tissues and two developmental time-points.

### Short sequence properties predictive of enhancers are conserved across species

We then investigated whether learned DNA sequence patterns predictive of enhancer activity were conserved across mammals by testing whether classifiers trained in one species could distinguish enhancers from the genomic background in another species. First, we applied the human liver classifier to the five other species. We quantified cross-species performance using the relative auROC—the auROC of the enhancer classifier trained on species A and applied to species B, divided by the average auROC obtained by the classifier trained and tested on species B. In other words, the relative auROC is the proportion of within-species performance achieved by a classifier trained in a different species. The classifier trained on human liver enhancers predicted liver enhancers in other mammals nearly as accurately as classifiers trained in each species (Figure 3a, PR curves in Figure S1c), and its relative performance decreased only slightly across species (Figure 3b, relative auROCs > 95.5%). Furthermore, the scores from the human classifier applied to human enhancers were significantly positively correlated with the scores from non-human classifiers (Figure S3; Spearman’s ρ between 0.90 for macaque and 0.66 for opossum). When expanded to all pair-wise combinations of species, classifiers accurately predicted enhancers in every mammalian species tested, regardless of the specific species they were trained in; the average relative auROC was 96.0% (Figure 3b; raw AUCs in Figure S4a-b).

**Fig 3.**
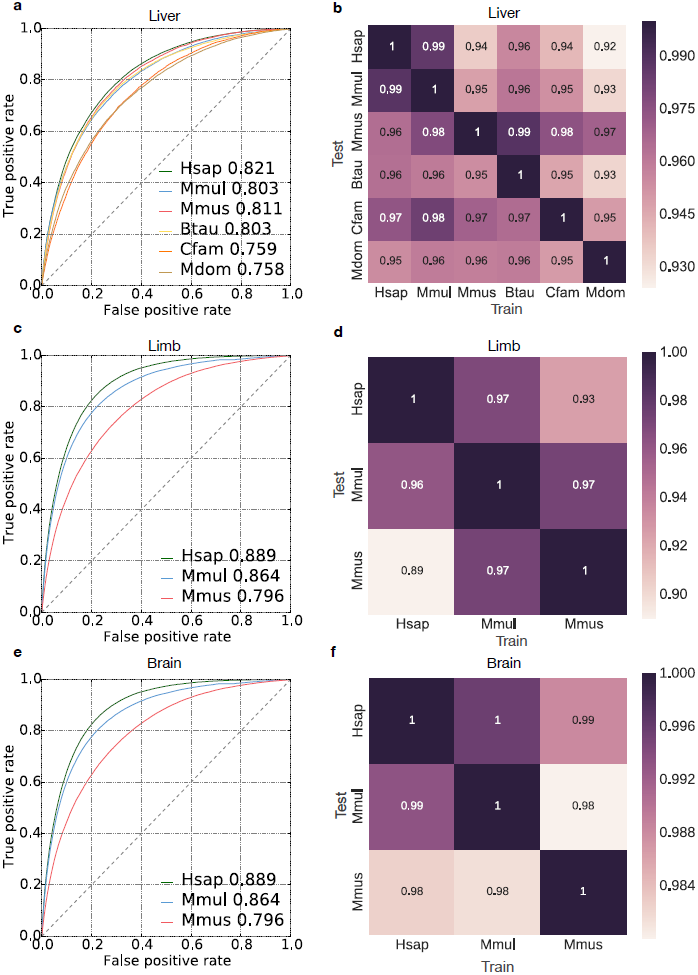
Human-trained enhancer classifiers accurately predicted liver, limb and brain enhancers in diverse mammals. (a) ROC curves of the performance of the human liver enhancer classifier applied to the human (Hsap), macaque (Mmul), mouse (Mmus), cow (Btau), dog (Cfam) and opossum (Mdom) datasets. Area under the curve (auROC) values are given after the species name. (b) Heat map showing the relative auROC of liver enhancer classifiers applied across species compared to the performance of classifiers trained and evaluated on the same species (Figure 2a). The classifiers were trained on the species listed on the x-axis and tested on species on the y-axis. (c) ROC curves showing the performance of the human limb enhancer classifier on human, macaque and mouse. (d) Heat map showing the relative auROC of limb enhancer classifiers applied across species compared to the performance of classifiers trained and evaluated on the same species (Figure 2b). (c) ROC curves showing the performance of the human brain enhancer classifier on human, macaque and mouse. (d) Heat map showing the relative auROC of brain enhancer classifiers applied across species compared to the performance of classifiers trained and evaluated on the same species (Figure 2c). The raw auROC and auPR values for all comparisons are given in Figure S4, Figure S6 and Figure S7.

Classifiers generalized better to more closely related species; generalization was significantly inversely correlated with the species’ evolutionary divergence, as quantified by substitutions per neutrally evolving site (Figure S5, Pearson’s r = –0.585, *P* = 0.022). Furthermore, classifiers trained to identify enhancers in developing limb and brain also accurately generalized across species. The average relative auROC for the developing limb classifiers was 95.0% across all species pairs (Figure 3c-d, Figure S6), and the average relative auROC for the developing brain classifiers was 98.6% (Figure 3e-f; raw AUCs in Figure S7). The ability of classifiers to generalize to other species illustrates the conservation of sequence properties predictive of enhancers across mammals.

To ensure that the small fraction of liver enhancers shared between pairs of species were not driving performance, we identified enhancers human liver enhancers that overlapped enhancers from each of the other five mammalian species in genome-wide multiple sequence alignments from Ensembl (macaque: 24.0%; mouse: 13.6%; cow: 20.0%; dog: 16.7%; and opossum: 3.4%). For each pair of species, the overlapping enhancers were removed from the human training set and a new human classifier was trained and evaluated. Removal of these enhancers had little impact on classifier performance across species, illustrating that the small fraction of orthologous enhancers did not drive the cross-species generalization of the classifiers (Figure S8).

Genome-wide mapping of enhancer-associated histone modifications is a cost-effective means to identify putative enhancers; however, the presence of these modifications does not guarantee enhancer activity. Many experimental and computational approaches have been used to identify enhancers [1,49], and there is considerable disagreement among different strategies [50]. To investigate the generality of our conclusions drawn from histone-modification-derived enhancers, we also analyzed enhancers validated *in vivo* via transgenic assays from the VISTA enhancer database. We included six tissues (limb, forebrain, midbrain, hindbrain, heart and branchial arch) with a sufficient number of validated enhancers (at least 50) in human and mouse. Consistent with the results from classifiers trained on histone-modification defined enhancers, the classifiers trained and evaluated on VISTA human enhancers accurately predicted VISTA mouse enhancers in the corresponding tissue from genomic background, and vice versa (Figure S9, average relative auROC = 96.3%). This suggests that sequence patterns in enhancers confirmed via reporter assays are conserved between human and mouse. Moreover, the histone-modification trained limb classifiers accurately predicted VISTA enhancers (auROC = 0.83 in human, 0.76 in mouse) competitively with the VISTA-trained limb classifier itself (auROC = 0.81 in human, 0.78 in mouse), suggesting that sequence properties predictive of histone-modification defined enhancers are also predictive of transgenic assay validated enhancers. Thus, in spite of the limited number and biases present in the sequences tested for enhancer activity by VISTA, these analyses demonstrate that our models capture conserved sequence attributes of functionally validated enhancers.

Overall, these results show that the DNA sequence profiles of enhancer sequences captured by species-specific 5-mer spectrum SVM classifiers are predictive of enhancers in other mammalian species in corresponding tissues. The strong generalization of performance and correlation of the predictions for specific sequences by classifiers trained in different species indicates that sequence properties predictive of enhancers are conserved across mammals.

### Short DNA sequence patterns remain predictive of enhancer activity after controlling for GC content and repetitive elements

Enhancer activity is positively correlated with GC content (Figure S3), and enhancers can be born from repetitive sequences derived from transposable elements [51–54]. Thus, we sought to evaluate the extent to which these properties influenced the generalization of our enhancer prediction models across species. First, we trained GC-controlled classifiers using negative sets of random genomic regions matched on GC content. The predictive power of the GC-controlled classifiers was substantial (average auROC of 0.75 for liver, 0.79 for limb and 0.81 for brain; Figures S10a, S11a and S12a), but as expected, less than the corresponding classifiers without GC-control (average auROC of 0.81 for liver, 0.89 for limb and 0.92 for brain; Figure 2). Nevertheless, GC-controlled classifiers maintained strong cross-species generalization: liver classifiers had an average relative auROC of 94.8% when applied to the other five species (Figures 4a); limb classifiers had an average relative auROC of 95.0% when applied across species (Figures S11e); brain classifier had an average relative auROC of 94.8% (Figure S12e). The enhancer predictions for individual sequences by the GC-controlled classifiers were significantly correlated, and as expected, high GC-content sequences no longer received consistently high scores (Figure S13). Ultimately, the strong cross-species generalization of the GC-controlled classifiers suggests that enhancers differ from the genomic background in higher order sequence patterns beyond GC-content, and that those patterns are conserved.

The generalization of each liver GC-controlled classifier across species had the same pattern as the classifiers without GC-control: the human classifier had the best generalization (average relative auROC = 96.1%), while the opossum had the worst (average relative auROC = 92.8%). In these GC-controlled analyses, we observed a stronger inverse correlation between the relative performance across species and sequence divergence (Figure S14, Pearson’s *r* = –0.77, *P* = 0.001) than in the non-GC-controlled analysis (Figure S4, Pearson’s *r* = –0.585, *P* = 0.022). This indicates that both genomic differences in GC content distribution and overall evolutionary divergence influence the conservation of the sequence patterns predictive of putative enhancers.

To evaluate the influence of repetitive elements on the ability to distinguish enhancers from the background and the observed conservation of sequence properties across species, we trained classifiers to distinguish enhancers that did not overlap a repetitive element (only 3.3% of all enhancers in human) from matched non-repetitive regions from the genomic background. Neither the ability to distinguish enhancers from the background in a species, nor the ability of predictive sequence properties to generalize across species, was substantially reduced (Figure S15). This demonstrates that, while repetitive elements contribute to enhancer activity, the conservation of sequence properties predictive of enhancers is not contingent on their presence.

To examine the influence of repetitive elements across all observed enhancer sequences, we also trained classifiers to distinguish all enhancers regions from genomic background regions matched for both GC-content and the proportion of overlap with a repeat element. The performance of these classifiers slightly decreased (average auROC of 0.73; Figure S16a) relative to when not controlling for repeat overlap (average auROC of 0.75; Figure S10a) or neither repeats or GC-content (average auROC of 0.81; Figure 2). This indicates that, as expected, both features are informative about enhancer function. However, the repeat and GC-controlled classifiers still generalized across species (average relative auROC = 94.0%, Figure 4b); this demonstrates that enhancer sequence properties beyond both GC and repeat content are conserved across species.

**Fig 4.**
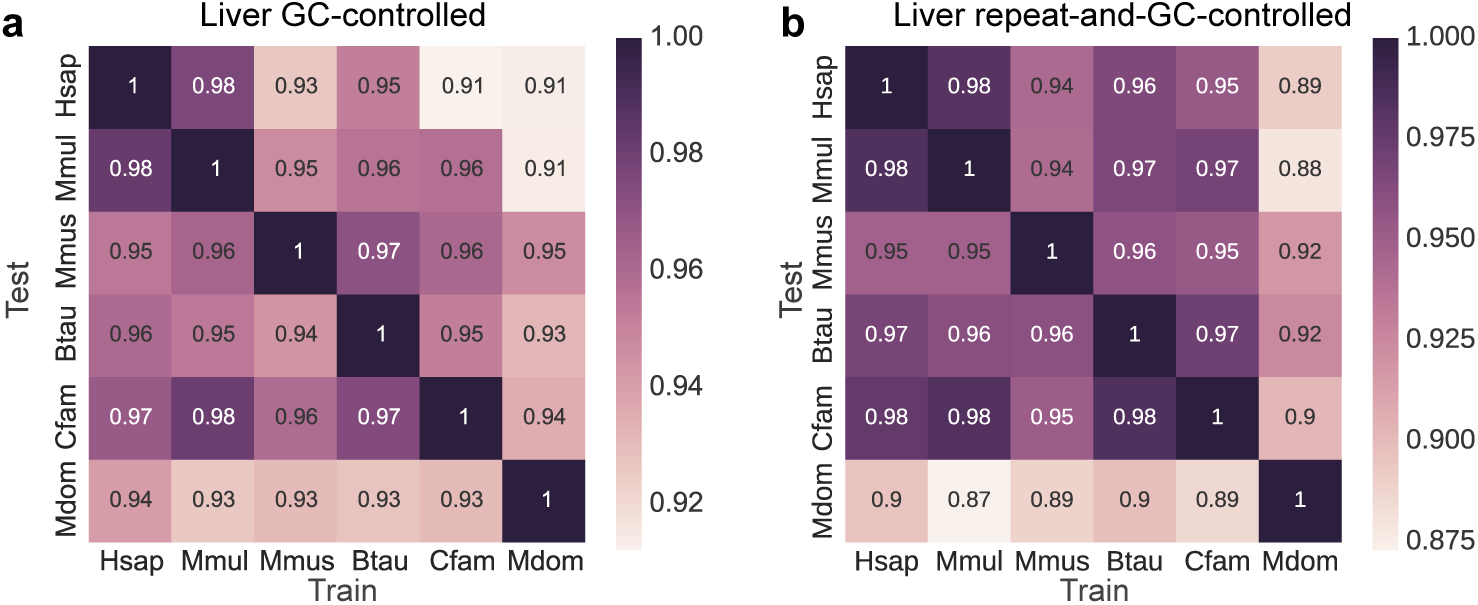
Enhancer sequence properties remain conserved across diverse mammals after controlling for both GC-content and repetitive elements. The heat maps give the cross-species relative auROCs for SVM classifiers trained on 5-mer spectra to identify enhancers in the species along the x-axis, and then used to predict enhancers in the species on the y-axis. The “negative” training regions from the genomic background were matched to the enhancers’: (a) GC-content, and (b) GC-content and proportion overlap with repetitive elements.

### Enhancer sequence properties are more similar across the same tissue in different species than across different tissues in the same species

Gene expression patterns are significantly more similar in corresponding tissues across species than between different tissues in the same species [14–16], and we demonstrated that enhancer sequence properties are strongly conserved in the same tissue across species (Figure 2). Thus, we hypothesized that, as for gene expression, enhancer sequence properties would be more similar in the same tissue across species (cross-species) than between different tissues in the same species (cross-tissue) and that the cross-tissue performance could provide a benchmark for contextualizing cross-species generalization. To test this, we performed cross-tissue analysis using human enhancers identified in nine diverse cellular contexts, including liver, by the Roadmap Epigenomics Project [2] (Methods). We applied the classifier trained on human liver enhancers (from Villar et al.) to Roadmap Epigenomics enhancers from: liver, brain hippocampus middle, pancreas, gastric, left ventricle, lung, ovary, CD14 cells, and bone marrow. For consistency, we used H3K27ac without H3K4me3 to identify enhancers in these tissues. Next, we compared the relative auROC between the cross-tissue and cross-species prediction tasks (Figure 5a). In the non-GC-controlled analysis, the human liver enhancer classifier predicts enhancers in macaque, mouse, cow, dog and opossum better than all non-liver Roadmap tissues. In the GC-controlled analysis, we observed the same trend. The cross-species predictions are also more accurate than cross-tissue predictions, with the exception of the Roadmap gastric tissue (dark green), which is also a digestive tissue. When compared to the relative auROCs of all pairwise cross-species analysis in liver, limb and brain, those of human liver to non-liver Roadmap tissues are significantly lower (Figure 5b). In addition to the human cross-tissue analysis, we also examined the cross-tissue performance of the liver, limb and brain classifiers over all three species with enhancers in three tissues: human, macaque and mouse. For each species, we applied the classifiers trained in liver, limb and brain to that species’ enhancers in other two tissues. Again, cross-species performance (all pairwise relative auROCs) was significantly higher than cross-tissue performance in both GC-controlled and non-GC-controlled analyses (Figure 5b). The ability of enhancers to regulate gene expression is often contingent on both cell-type specific attributes, such as expression patterns of TFs [55], and properties that are shared across active enhancers in general. The stronger performance of the trained classifiers in the cross-species compared to cross-tissue prediction tasks suggests that they capture cell-type-specific sequence attributes and that these features are conserved across species.

**Fig 5.**
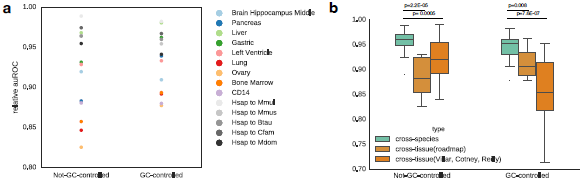
Enhancer classifiers generalize more accurately across the same tissue in different species than across different tissues in the same species. (a) The human-trained liver classifier obtains better performance when applied to liver enhancers from other species (gray dots) than when applied to enhancers from other human tissues. This also holds for GC-controlled analyses, with the exception of predicting enhancers active in the gastric mucosa. (b) In the not-GC-controlled analysis, the cross-species performance (average relative auROC = 96.2%) is significantly better than the cross-tissue (roadmap) performance (88.4%, Mann Whitney U test, *P* = 0.00005) and the cross-tissue (Villar, Cotney, Reilly) performance (92.0%, Mann Whitney U test, *P* = 2.2E-05). This also holds true for the GC-controlled analysis. The cross-species performance (average relative auROC = 94.6%) is significantly better than the cross-tissue (roadmap) performance (91.2%, Mann Whitney U test, *P* = 0.008) and the cross-tissue (Villar, Cotney, Reilly) performance (85.8%, Mann Whitney U test, *P* = 7.6E-07).

### The most predictive sequence patterns in different species match binding motifs for many of the same transcription factors

To interpret the biological relevance of the sequence patterns learned by the trained SVM enhancer prediction models in each species, we analyzed the similarity of the sequence properties in their functional context: TF binding motifs. First, we matched the 5% (n = 52) most enhancer-associated 5-mers learned by the human GC-controlled liver classifier to a database of 205 known TF motifs [56] using TOMTOM (Figure 6a). The enhancer-associated 5-mers were significantly more likely to match at least one TF motif than expected at random (46.1% vs. 27.7%; one-tailed *P* = 0.0035, binomial test). The 5% (n=52) most background-associated 5-mers were not significantly different from random (21.6% matched at least one TF, two-tailed *P* = 0.43, binomial test). This illustrates that the classifiers learned sequence patterns with regulatory potential.

**Fig 6.**
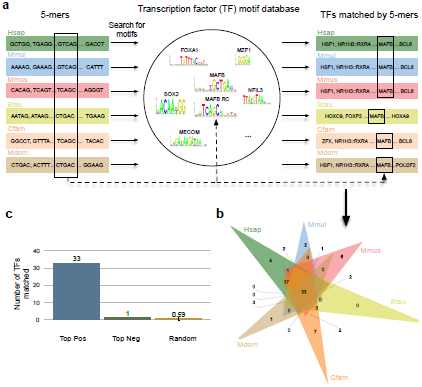
The DNA sequence patterns most predictive of liver activity across species matched a common set of transcription factors. (a) Transcription factor analysis workflow. For each species enhancer classifier, we found TF motifs matched by the top 5% positively weighted 5-mers. Note that different 5-mers (marked with black box on the left) can match the same motif, e.g., MAFB and its reverse complement (RC). The overlap of matched TFs were then compared across each species’ classifier. (b) Venn diagram of the sharing of the TF motifs matched by the top 5% positive 5-mers from each GC-controlled liver classifier. The total number of TFs matched by top 5-mers in each species was: 121 (human), 104 (macaque), 100 (mouse), 81 (cow), 118 (dog), 102 (opossum). Similar results were observed for the non-GC-controlled classifier (Figure S17a). (c) The number of TFs matched by all species based on 5-mers in top positive, top negative, and 100 random sets of 5% of all possible 5-mers. The 33 TF motifs shared among the high-weight set for each species is thus significantly more than expected.

Next, we investigated whether the TF binding motifs matched by enhancer-associated 5-mers were shared between species. The highly weighted 5-mers in the human-trained classifier matched 121 TF motifs. Of these, the binding motifs for 33 TF were also matched by enhancer-associated 5-mers in all other species (Figure 6b, Table S1). This is significant enrichment for shared TF motifs among the enhancer-associated 5-mers; only 0.59 TF motifs were shared across all species on average over 100 random sets of 5% of 5-mers from each species (Figure 6c). Similarly, only one TF motif (MZF1) was shared among all the species’ most background-associated 5-mers. The GC-controlled limb and brain classifiers also shared more TFs among the top 5% of enhancer-associated 5-mers than expected from random sets: 12 TFs were shared among the limb classifiers and 22 were shared among the brain classifiers vs. 8.1 shared TFs expected by chance. However, it is likely that the smaller number of available species for developing limb and brain enhancers, our limited knowledge of binding motifs for TFs active in developing limb and brain, and the heterogeneity of developing limb and brain tissue reduced power to detect sharing compared to liver. We obtained similar results when comparing the TFs matched by 5-mers from non-GC-controlled SVM models (Figure S17).

To evaluate the relevance of the shared TF motifs to liver function, we evaluated the expression patterns of the TFs across 12 tissues [57]. Shared TFs among liver enhancer-associated 5-mers were significantly enriched for liver expression (Table 1, *P* = 0.011, one-tailed Fisher’s exact test). Many of the shared TFs play an essential role in liver function. For instance, they are enriched for activity in the TGF-β signaling pathway compared to non-shared TFs; the enrichment is mainly due to members of the AP-1 (JUN, FOS, and MAF subfamilies) and SMAD families (Methods) [58,59]. TGF-4 signaling is a central regulatory mechanism that is disrupted in all stages of chronic liver disease [60]. Further, mice deficient in c-JUN or MAF have an embryonic lethal liver phenotype [61,62]. We also searched for matches to the binding motifs of known liver master regulators among the highly weighted motifs. While none of them were shared among all models, several including, HNF1α, HNF4α, and FOXA1 matched highly weighted motifs in three or more species (Table S2). This demonstrates that the sequence patterns learned in each species capture similar motifs that are recognized by TFs that important to the relevant tissue context.

**Table 1.**
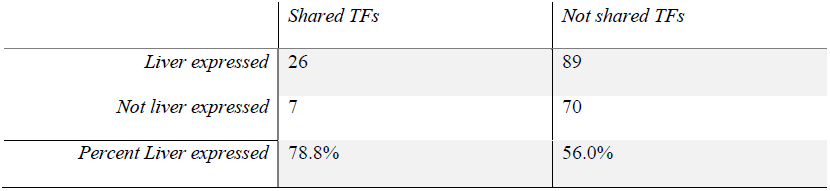
The TFs with motifs shared among the top 5-mers across all species’ liver enhancer SVM classifiers are significantly enriched for liver expression (*P* = 0.011, one-tailed Fisher’s exact test).

|  | *Shared TFs* | *Not shared TFs* |
| --- | --- | --- |
| *Liver expressed* | 26 | 89 |
| *Not liver expressed* | 7 | 70 |
| *Percent Liver expressed* | 78.8% | 56.0% |

### Convolutional neural networks predict enhancers more accurately than SVMs, but generalize less well across species

Although the *k-*mer SVM model accurately classified many enhancers, its performance was not perfect. We hypothesized that using models with the potential to learn combinatorial interactions between short sequence patterns in enhancers could further improve performance. To model these more complex patterns, we trained convolutional neural networks (CNNs) to distinguish liver enhancers from the genomic background in each species. Here, we used a balanced dataset due to challenges of training CNN classifiers on unbalanced sets (Methods). To compare the performance of CNNs with the SVM models, we retrained *k*-mer spectrum SVM classifiers on the same balanced data in each species and performed cross-species enhancer predictions (Supplementary Figure 18a,b). The CNN model performance is substantially better than the *k*-mer spectrum SVM classifiers at predicting enhancers from genomic background in each species (Figure 7a), suggesting that the ability to model complex interactions between short sequence patterns improves predictions. Moreover, the first layer of the human liver CNN learned many binding motifs for TFs relevant to liver biology, including CEBPB, HNF4A, and HNF1A (Figure 7b).

**Fig 7.**
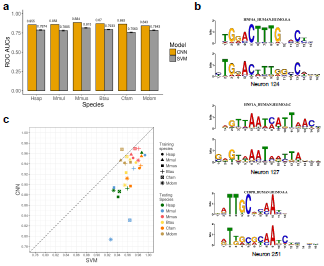
CNNs identify enhancers more accurately than 6-mer-based SVM models, but generalize less well across species. (a) The auROCs of CNN models perform substantially better than the 6-mer SVM model in each species. The error bars give the standard error of ten-fold cross-validation for the SVM models. (b) Neurons in the first layer of the CNN learned the motifs of important liver TFs, including HNF4A, HNF1A, and CEBPB. (c) The relative auROCs of the CNN models applied across species are consistently lower than for the 6-mer SVMs applied across the same species. This suggests that the CNN models do not generalize as well across species as the SVM models.

Next, we performed the cross-species enhancer prediction with the CNNs. The CNN models generalize well across species (relative auROC from 0.79 to 0.97), but their generalization is consistently worse than the *k*-mer SVM models (Figure 7c; Raw auROCs and auPRs is in Supplementary Figure 18c,d). This suggests that the combinatorial sequence patterns captured by the internal layers of CNN models is less conserved across species then the individual short sequence motifs.

## Discussion

In this study, we trained SVM and CNN classifiers based on DNA sequence patterns to distinguish enhancers from the genomic background in diverse mammalian species. We then showed that, in spite of significant changes in the enhancer landscape between species, the *k*-mer SVM models trained using short sequence patterns as features exhibited minimal decreases in performance when applied across species. This indicates that short sequence patterns predictive of enhancer activity captured by these models are conserved across mammals. Furthermore, the DNA patterns most predictive of activity across species matched a common set of TF binding motifs with enrichment for expression in the relevant tissues. The sequence properties predictive of histone-mark defined enhancers were also predictive of enhancers confirmed in transgenic reporter assays. We then showed that CNN models performed better than the SVMs at identifying enhancers, but they generalized less well across species. These results suggest that conserved regulatory mechanisms have maintained constraints on short sequence motifs present in enhancers for more than 180 million years, while more evolutionary change in regulatory mechanisms has occurred at the level of combinations of motifs, likely representing cooperative interactions between TFs.

Confidently identifying and experimentally validating enhancers remains challenging [50]. We showed that short sequence properties are conserved across species using enhancers identified via two complementary techniques: histone modification profiling and transgenic assays. Each of these approaches has strengths and weaknesses. The histone modification based enhancer predictions enable genome-wide characterizations across many species, but this approach is prone to false positives. On the other hand, the transgenic assays clearly demonstrate the competence of a sequence to drive gene expression, but are restricted to a biased set of relatively few sequences from two species that are tested at one developmental stage. By showing the cross-species conservation is maintained in both categories, and that models trained on each set perform similarly, we argue the conservation of enhancer short sequence properties is robustness to the methodology used to define enhancers.

The design of this study can serve as a framework for further examining the conservation and divergence of regulatory sequence patterns across species. We trained sequence-based machine learning models within a species, and then applied them to other species; this approach can be applied on a genome-wide scale, is not dependent on knowledge of TF binding motifs, and allows some flexibility in the weights assigned to each feature while directly testing the generalization of overall sequence patterns. Identification of enhancers in more divergent species would enable us better quantify how deeply conserved enhancer sequence properties are. This remains an open question, as more divergent animal species have very little conservation of TF co-associations at putative enhancers despite conservation of TF binding preferences [63]; however, enhancer properties appear to be conserved over greater evolutionary timescales in insects [41,42,64]. Identification of enhancers in the same cellular context for more closely related species would also be valuable to enable the investigation of lineage-specific regulatory sequence patterns. Thus, additional comparative studies of regulatory sequence features in more species are needed to better understand both recent and ancient influences on regulatory sequences.

While both the SVM and CNN classifiers correctly distinguished many enhancers from the genomic background, neither performed perfectly. Many factors contribute to this, including: false positives in the training data, noise from the low resolution of the histone modification peaks (i.e., they include non-functional sequence flanking the enhancer), and the features considered in our models. As enhancer datasets and prediction methods improve, it will be valuable to continue to evaluate generalization across species. Additionally, the features learned by the enhancer CNNs are difficult to interpret biologically, especially for higher-level neurons. The interpretation of internal layers of accurate CNNs would facilitate the understanding of how more complex rules of the enhancer sequence architecture, such as motif spacing, order, combinations, and hierarchies, change during evolution. The interpretation and comparison of conserved and diverged rules between species is an important area for future work. Furthermore, our framework could also be adapted to investigate conservation of other functionally relevant factors, such as histone modifications and DNA shape [10,65].

## Conclusions

We demonstrated that short DNA sequence patterns predictive of enhancer activity learned in one species generalize very well to other mammals. Furthermore, deep neutral network models that can learn complex combinations of short sequence patterns identified enhancers even more accurately, but generalized less well across species. This suggests evolutionary conservation short sequence motifs, but turnover of their some of their combinatorial patterns between species. The commonality of short sequence elements predictive of enhancer activity across mammals argues that much of what we learn about enhancer biology, particularly at the basic sequence motif level, in model organisms could be extrapolated to humans. Sequence-based cross-species enhancer prediction could be of particular use in studying difficult to obtain human tissues and providing preliminary annotations in uncharacterized species and tissues. There is also the potential to combine sequence-based models with successful cross-species enhancer prediction strategies based on functional genomics data [66]. Nonetheless, much work remains to understand how regulatory programs are robust to sequence changes, yet receptive to functional divergence, and to facilitate our interpretation of the effects of non-coding variants in diverse mammals.

## Methods

### Genomic data

All work presented in this paper is based on hg19, rheMac2, mm10 (mouse liver dataset), mm9 (mouse limb and brain dataset), bosTau6, canFam3 and monDom5 DNA sequence data from the UCSC Genome Browser. For consistency with the original studies, liver gene annotations are from Ensembl v73, limb and brain gene annotations are from Ensembl v67 [67]. The sequence divergence between each pair of species was computed from the neutral model built from fourfold degenerate sites in the 100-way multiple species alignment from UCSC Genome Browser (http://hgdownload.cse.ucsc.edu/goldenpath/hg19/phastCons100way/).

### Enhancer and genomic background datasets

We evaluated the ability of machine learning models to distinguish different sets of enhancers (positives) from sets of matched regions from the genomic background (negatives). In this section, we describe the collection and processing of the enhancer and genomic background sets. In the next section, we describe the training and evaluation of the SVM classifiers.

We analyzed three multi-species histone-modification-defined enhancer datasets in this study. The first consisted of liver enhancers identified by genome-wide ChIP-seq profiling of histone modifications (H3K27ac without H3K4me3) in 20 species from five mammalian orders [12]. We selected a member of each order with a high-quality genome build for analysis when possible; however, the most diverged order—marsupials—did not have a species with a high-quality genome build. We consequently selected opossum, as it was the most diverged from humans. This resulted in the following species, with the number of observed enhancers in each: human (N=29512), macaque (N=22911), mouse (N=18517), cow (N=30892), dog (N=18966), and opossum (N=23160) [12].

We generated three different sets of matched genomic background regions for use as negatives in the training and evaluation of the liver classifiers for each of the six species. The first are random genomic regions matched on length and chromosome to the observed enhancers. Second, for the GC-controlled analyses, we generated genomic background regions matched to the enhancers on length, chromosome, and GC-content. Finally, for the repeat controlled analysis, we obtained repetitive elements identified by RepeatMasker for each species [68] and generated random regions from the genomic background matched on length, chromosome, GC-content, and proportion overlap with repetitive elements. To reflect the fact that enhancers make up a small portion of the genome, we chose an imbalanced data design with 10 times as many of the genomic background (negative) regions as there were enhancers. For all analyses, we did not consider enhancers or random regions that fell in genome assembly gaps (UCSC gap track) when generating negatives. For human and mouse, we also excluded the ENCODE blacklist regions [57] https://sites.google.com/site/anshulkundaje/projects/blacklists).

The second enhancer dataset contained human (N=25304), macaque (N=88560), and mouse (N=87406) enhancers identified from profiling the H3K27ac modification in developing limb tissue [8]. The third enahncer dataset contained human (N=48853), macaque (N=57446), and mouse (N=51888) enhancers identified from profiling the H3K27ac modification in developing brain tissue [69]. For limb and brain enhancers, we excluded regions within 1 kb of a transcription start site. For each species, we combined the enhancer regions from different development stages. The genomic background regions for each species were defined following the same procedure as for the liver enhancers.

To determine how well classifiers generalized across additional tissue types, we used human enhancers identified by the Roadmap Epigenomics Project [2] in nine tissues from diverse body systems: liver (GI, E066), hippocampus middle (brain, E071), pancreas (exocrine-endocrine, E098), gastric (GI, E094), left ventricle (heart, E095), lung (E096), ovary (reproductive, E097), bone marrow derived mesenchymal stem cell cultured cells (stromal-connective, E026) and CD14 primary cells (white blood, E029). We defined enhancers in these tissues as H3K27ac without H3K4me3 regions. For each tissue, we generated not-GC-controlled and GC-controlled negative training examples as described for the liver enhancers above.

In addition to the histone-modification-defined enhancers, we also analyzed enhancers validated in transgenic reporter assays in embryonic day 11.5 mouse embryos from VISTA [70]. We investigated all six tissues with at least 50 positive enhancer elements in both species: forebrain, midbrain, hindbrain, limb, heart and branchial arch. These enhancers comprised the positive training examples. For each tissue, we additionally generated 10 times the number of enhancers length and chromosome matched random genomic regions as negative training examples.

### Spectrum kernel SVM classification

An SVM is a discriminative classifier that learns a hyperplane to separate the positive and negative training data in feature space. We used the *k*-mer spectrum kernel to quantify sequence features for the SVM [71]. Training, classification, evaluation, and the computation of features weights were performed with the kebabs R package (v1.4.1) [48]. We used the default kernel normalization to the unit sphere, considered reverse complements separately, used the cosine similarity, and used a cost parameter (*C*) of 15. Due to the imbalanced training dataset, we set class weights of 10 for the positives and 1 for the negatives to increase the penalty on misclassification of positives. We report all analyses with *k* = 5, but classifier performance and generalization were similar for *k* = 4–7 (0.81, 0.82, 0.82, 0.82, respectively for liver).

To evaluate classifier performance within-species, we performed ten-fold cross validation. In other words, for each set of positives and negatives, the entire data set was randomly partitioned into ten independent sets that maintained the ratio of positives and negatives. Positives and negatives from nine of the ten sets were then used to train the classifier, the trained classifier was then applied to the remaining partition, and these predictions were used to evaluate the classifier. This process was performed ten times, testing each partition once. To summarize performance, we averaged the auROC and auPR over the ten runs. For cross-species classification, we trained on the whole dataset in the training species and evaluated the performance on a random half of the dataset in the test species due to computational limitations of the kebabs package.

We also evaluated more flexible models, such as the mismatch [48,71] and gappy pair kernels [48,72], These *k*-mer-based prediction models are similar to the spectrum kernel, but the mismatch kernel allows a maximum mismatch of *m* nucleotides in the *k*-mer and the gappy pair kernel considers pairs of *k*-mers with maximum gap of length *m* between them. For comparison, we trained the gappy pair kernel with *k* = 2, *m* = 1 and mismatch kernel with *k* = 5, *m* = 1 to compare with the 5-mer spectrum kernel. The mismatch and gappy pair kernels did not significantly increase the performance (auROCs of 0.82 and 0.82, respectively for liver) and are less interpretable than the *k*-mer spectra (Figure S2). It is possible that other parameter settings could yield slightly improved performance, but the resulting models would be more difficult to interpret, and optimizing performance was not the goal of our study.

### Transcription factor motif analysis

5-mers were matched to known TF binding motifs in the JASPAR 2014 Core vertebrate database [56] using the TOMTOM package with default parameters [73]. The sharing of 5-mers and TFs across species was visualized using jVenn [74].

### Transcription factor expression data

We obtained RNA-seq data for TFs across 12 tissues from the Gene Expression Atlas (https://expressionatlas.org/hg19/adult/). Genes with non-zero FPKM (Fragments Per Kilobase of transcript per Million mapped reads) in a tissue were considered as expressed.

### Convolutional neural network (CNN) classifiers and comparison to *k*-mer SVM models

We used the center 3000 bp (approximately the median length) of liver enhancers in six selected species as the positive training sequences and the same number of length matched random genomic regions in the corresponding species as negative training sequences. We split the dataset into training (80%), validation (10%), and a hold-out test set (10%).

A typical convolutional neural network consists of convolutional layers, max-pooling layers, fully connected layers, and an output layer. To define the CNN structure (Figure S19), we first defined a hyperparameter space, including a range of learning rates, number of layers, the window size of the filters (neurons), and regularization strength. Next, we trained 100 human CNN models to identify liver enhancers using keras [75] with hyperparameters suggested by the Tree-structured Parzen Estimator (TPE) approach implemented in the hyperopt [76] library and selected the best set of hyperparameters based on the smallest loss in the human validation set. Then, we trained the enhancer CNN model with the best human CNN structure in the other five species, but different regularization strengths 30 times in order to find the best performing CNN model for each species. The performance of within-species prediction is reported based on the auROC of predicting the hold-out set of the training species and the performance of cross-species prediction is reported based on the auROC of predicting all data in the testing species.

To interpret the first layer of the human liver CNN, we forward propagated sequences in the human liver validation dataset through the CNN and selected the sequence patches that maximally activate each neuron (> 0.5 maximum activation value of the neuron) in the first layer. Then, we converted the resulting sets of sequence patches to position weight matrices (PWMs) and mapped the PWMs to human TF motifs from the HOCOMOCO v11 [77] database using TOMTOM with default parameters [73].

For direct comparison to the performance of CNNs, we also trained *k*-mer spectrum SVM models for each species on the same balanced dataset at the CNNs. We compared the performance for *k* from 4 to 8 on this balanced human dataset; we report results for *k* of 6 based on it giving the best average auROC in ten-fold cross-validation. The performance of within-species prediction is reported based on the average auROC of ten-fold cross validation and the performance of cross-species prediction is reported based on the auROC of predicting all data in the testing species.

## Declarations

### Acknowledgements

We thank D. Kostka, D. Rinker, and C. Simonti for helpful discussions and comments on the manuscript.

### Availability of data and materials

The source code for cross-species enhancer prediction and generating all results during the current study is available at: https://github.com/lingchen42/EnhancerCodeConservation.

### Author Contributions

J.A.C. conceived and supervised the project. L.C. collected the data and led the analyses. A.E.F. contributed analyses. All authors interpreted the data and wrote the manuscript.

### Consent for publication

Not applicable.

### Ethics approval and consent to participate

Not applicable.

## Supporting information

**S1 Appendix. Supplementary figures S1–S19**

**S1 Table. Liver expression of the shared TF motifs in the liver GC-controlled analysis**

**S2 Table. The sharing of the TF motifs matched by the top 5% positive 5-mers from each classifier.**

